# Building realistic assemblages with a Joint Species Distribution Model

**DOI:** 10.1101/003947

**Authors:** David J. Harris

## Abstract

1. Species distribution models (SDMs) can be used to predict how individual species—and whole assemblages of species—will respond to a changing environment. Until now, these models have either assumed (1) that species’ occurrence probabilities are uncorrelated, or (2) that species respond linearly to preselected environmental variables. These two assumptions currently prevent ecologists from modeling assemblages with realistic co-occurrence and species richness properties.
2. This paper introduces a stochastic feedforward neural network, called mistnet, which makes neither assumption. Thus, unlike most SDMs, mistnet can account for non-independent co-occurrence patterns driven by unobserved environmental heterogeneity. And unlike recently proposed Joint SDMs, mistnet can also learn nonlinear functions relating species’ occurrence probabilities to environmental predictors.
3. Mistnet makes more accurate predictions about the North American bird communities found along Breeding Bird Survey transects than several alternative methods tested. In particular, typical assemblages held out of sample for validation were nearly 50,000 times more likely under the mistnet model than under independent combinations of single-species models.
4. Apart from improved accuracy, mistnet shows two other important benefits for ecological research and management. First: by analyzing co-occurrence data, mistnet can identify unmeasured—and perhaps unanticipated—environmental variables that drive species turnover. For example, mistnet identified a strong grassland/forest gradient, even though only temperature and precipitation were given as model inputs. Second: mistnet is able to take advantage of incomplete data sets to guide its predictions towards more realistic assemblages. For example, mistnet automatically adjusts its expectations to include more forest-associated species in response to a stray observation of a forest-dwelling warbler.

## Introduction

Programs for managing and understanding biodiversity each require information about where species occur and where they could occur. Statistical approaches to these questions, such as species distribution models (SDMs), are important because they can help us anticipate how beneficial species might fare—or how harmful species might spread—in scenarios that we cannot observe directly (Elith & Leathwick 2009). Modern SDMs need not assume that species respond to environmental variation in a pre-specified way (e.g. linearly or quadratically); relaxing this assumption has substantially improved our ability to make predictions about where species can occur (Elith *et al*. 2006).

Unfortunately, existing nonlinear approaches do not always answer the most pressing questions for ecologists. Ecologists are not only interested in individual species; we are also interested in learning about higher-level patterns, such as community structure, species richness, species turnover, and alternative stable states (Chase 2003). While SDMs are often combined (“stacked”) to generate assemblage-level predictions (Pellissier *et al*. 2013), doing so requires assuming that species’ occurrence probabilities are uncorrelated (Clark *et al*. 2013; Calabrese *et al*. 2014). As shown in more detail below, ignoring these correlations leads stacked models to predict incoherent jumbles of species rather than realistic assemblages (Clark *et al*. 2013). A major source of non-independence among species—which stacked SDMs ignore—is shared dependence on unobserved environmental factors (McInerny & Purves 2011; Figure 1; Calabrese *et al*. 2014). Given that most models only use climate variables as predictors (Austin & Van Niel 2011), the set of unobserved factors will usually include *all of ecology* apart from climatic influences. SDMs’ failure to model other ecological processes is thus widely considered to be a major omission from statistical ecology’s toolbox (Austin & Van Niel 2011; Guisan & Rahbek 2011; Kissling *et al*. 2012; Wisz *et al*. 2013; Clark *et al*. 2013).

**Figure 1.**
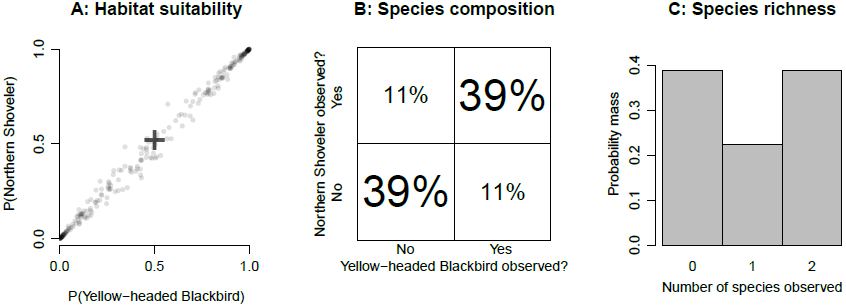
Unobserved environmental heterogeneity can induce correlations between species; ignoring this heterogeneity can produce misleading results. **A**: Based on climate predictors, a pair of single-species models might predict 50% occurrence probabilities for each of two wetland species (black cross). Climate predictors are not sufficient in this case, however: a site’s suitability for these species cannot really be determined without information about the availability of wetland habitat. Real habitats will to be tend to be suitable for both species (dense cloud of points in upper-right corner) or neither (lower-left corner), depending on this unmeasured variable. **B** This correlation among species substantially alters the set of assemblages one would expect to observe. (Under independence, all four possibilities would be equally probable.) **C P**ositive correlations among species can even induce a strongly bimodal distribution of species richness values.

In the last few years, several mixed models have been proposed to help explain the cooccurrence patterns that stacked SDMs ignore (Latimer *et al*. 2009; Ovaskainen, Hottola & Siitonen 2010; Golding 2013; Clark *et al*. 2013; Pollock *et al*. 2014). These *joint* species distribution models (JSDMs) can produce mixtures of possible species assemblages (points in Figure 1a), rather than relying on a small number of environmental measurements to fully describe each species’ probability of occurrence (which would collapse the distribution in Figure 1a to a single point; Pollock *et al*. 2014). In JSDMs (as in nature), a given set of temperature and precipitation measurements could be consistent with a number of very different possible sets of co-occurring species, depending on factors that ecologists have not necessarily measured or even identified as important. JSDMs represent these unobserved (latent) factors as random variables whose true values are unknown but whose existence would still help explain discrepancies between the data and the stacked SDMs’ predictions (Figures 1b and 1c). While JSDMs represent a major advance in community-level modeling (Clark *et al*. 2013; Pollock *et al*. 2014), existing implementations have all assumed that species’ responses to the environment are linear (in the sense of a generalized linear model). Thus, these JSDMs sacrifice the flexibility of modern single-species models, reducing their accuracy and limiting their utility.

Here, I present a new R package for assemblage-level modeling—called mistnet—that does not rely on independence (as stacks of single-species models do) or linearity (as previous JSDMs do). Mistnet is a stochastic feed-forward neural network (Neal 1992; Tang & Salakhutdinov 2013) that combines the nonlinear flexibility of modern single-species models with the latent variables found in previous JSDMs (cf Hutchinson, Liu & Dietterich 2011). In order to demonstrate the value of this approach, I compared mistnet’s predictive likelihood with that of several existing models, using observational data from thousands of North American Breeding Bird Survey transects (BBS; Sauer *et al*. 2011). A high predictive likelihood indicates that the model expects to see assemblages like those found along transects held out-of-sample, while a very low likelihood means that the model has effectively ruled those assemblages out due to overfitting or underfitting.

An accurate JSDM would up new possibilities for research and effective management. For example, although most models only have access to climate data (Austin & Van Niel 2011), a successful model of community structure should also be able to identify the major axes of non-climate variation that drive species turnover based on the species’ observed co-occurrence patterns. Moreover, a successful assemblage-level model would be able to take advantage of partially-completed samples or other kinds of prior information about a few species to inform its predictions about the rest of the assemblage. Since data collection efforts are frequently asymmetrical or incomplete, the ability to transfer information from well-documented taxa to more cryptic or rare species would prove valuable for community ecologists and conservationists alike. While a model’s ability to infer, for example, that “waterbirds like water” would not provide any novel biological insights, it would demonstrate that a modeling framework is ready to tackle more difficult problems where the biology is not already known.

## Materials and Methods Methods

Methods are presented in four main sections: (1) an introduction to the data sets used in this analysis, (2) a description of mistnet, (3) a summary of the existing methods used for model comparison, and (4) criteria for model evaluation.

## Data

Field survey data was obtained from the 2011 Breeding Bird Survey (BBS; Sauer *et al*. 2011). The BBS data consists of thousands of transects (“routes”), which I used as the main unit for my analysis. Each route includes 50 stops, about 0.8 km apart. At each stop, all the birds observed in a 3-minute period are recorded, using a standardized procedure. Following BBS recommendations, I omitted nonstandard routes and data collected on days with bad weather.

In order to evaluate SDMs’ capacities for predicting species composition, I split the routes into a “training” data set consisting of 1559 routes and a “test” data set consisting of 280 routes (Figure 2; Appendix A). The two data sets were separated by a 150-km buffer to ensure that models could not rely on spatial autocorrelation to make accurate predictions about the test set (Bahn & McGill 2007) (Appendix A). Each model was fit to the same training set, and then its performance was evaluated out-of-sample on the test set.

**Figure 2.**
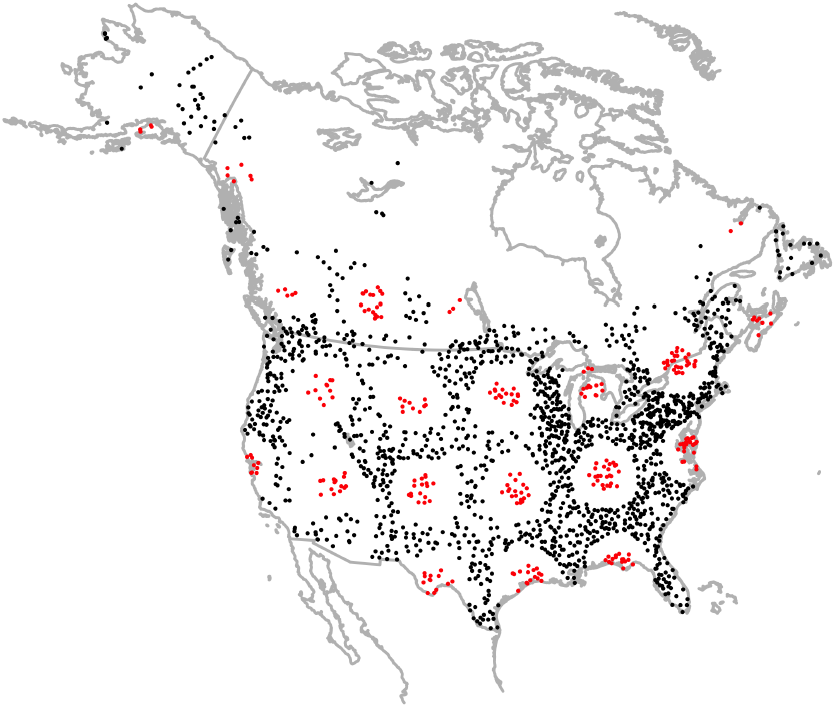
Map of the BBS routes used in this analysis. Black points are training routes; red ones are test routes. The training and test routes are separated by a 150-km buffer in order to minimize spatial autocorrelation across the two partitions.

Observational data for each species was reduced to “presence” or “absence” at the route level, ignoring the possibility of observation error for the reasons outlined in (Welsh, Lindenmayer & Donnelly 2013). It would be possible to incorporate the possibility of such errors in the model-fitting procedure if appropriate data were available, as was done in (Hutchinson *et al*. 2011). 368 species were chosen for analysis according to a procedure described in Appendix A.

To obtain environmental predictors for the model, I extracted the 18 Bioclim climate variables for each route from Worldclim (version 1.4; Hijmans *et al*. 2005). I omitted variables that were nearly collinear with one another (i.e. |*r*| >0.8) using the `findCorrelation` function in the `caret` package (Wing *et al*. 2013), leaving eight climate-based predictors (Appendix A). Since most SDMs do not use land cover data (Austin & Van Niel 2011) and one of mistnet’s goals is to make inferences about unobserved environmental variation, no other variables were included in this analysis.

Finally, I obtained habitat classifications for each species from the Cornell Lab of Ornithology’s All About Birds website (www.allaboutbirds.org) using an R script written by K. E. Dybala.

### Introduction to stochastic neural networks

Neural networks describe nonlinear mappings from input variables to predictions about one or more output variables. In general, ecologists have not had much success using neural networks for SDM, compared with other methods (e.g. Dormann *et al*. 2008). However, modern neural networks have recently outperformed other machine learning techniques in a wide range of applied contexts (Bengio 2013) and are thus worth a second look.

Mistnet models are *stochastic* neural networks, meaning that they include latent random variables (Neal 1992; Tang & Salakhutdinov 2013). In such a model, species’ occurrence probabilities are not fully specified the variables ecologists happen to measure, but can also depend on factors that have not been observed. In the absence of any information about these variables, mistnet (like other JSDMs) represents them using standard normal distributions. Depending on which values are sampled from these normal distributions and fed through the neural network, the model will expect to see different kinds of species assemblages (Figure 3).

**Figure 3.**
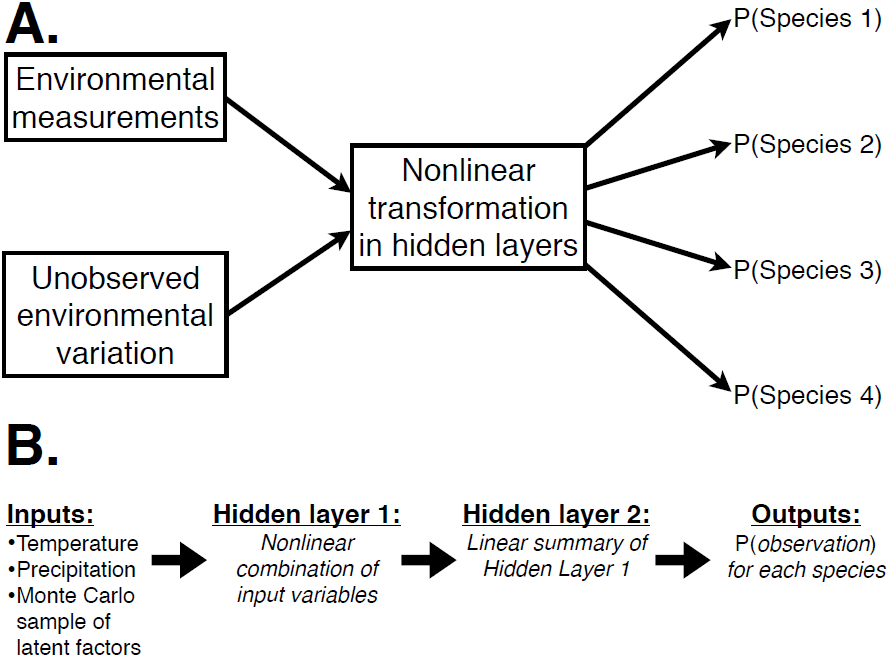
**A** A generalized diagram for stochastic feed-forward neural networks that transform environmental variables into occurrence probabilities multiple species. The network’s hidden layers perform a nonlinear transformation of the observed and unobserved (“latent”) environmental variables; each species’ occurrence probability then depends on the state of the final hidden layer. **B** The specific network used in this paper, with two hidden layers. The inputs include Worldclim variables involving temperature and precipitation, as well as random draws from each of the latent environmental factors. These inputs are multiplied by a coefficient matrix and then nonlinearly transformed in the first hidden layer. The second hidden layer uses a different coefficient matrix to linearly transform its inputs down to a smaller number of variables (like Principal Components Analysis of the previous layer’s activations). A third matrix of coefficients links each species’ occurrence probability to each of the variables in this linear summary (like one instance of logistic regression for each species). The coefficients are all learned using a variant of the backpropagation algorithm.

While the model’s main function is to make predictions about the species found in a given environment, inference can also proceed backward through the network, so that the presence (or absence) of a particular species can provide indications about the local environment—and thus about the likely configuration of the rest of the assemblage. This kind of inference could be useful in a variety of important contexts. For example, data is often more plentiful about waterfowl than about other wetland species, due to interest from hunters and conservation groups. If waterfowl are known to be present along a route, then a JSDM should recognize that suitable habitat was available, automatically increasing the estimated probability of occurrence for other species known to have similar habitat requirements. Notably, none of this extra inferential power requires that the mistnet user understand *which* environmental factors are driving the correlations between species, since these correlations are automatically inferred from species’ co-occurrence patterns.

The neural network used here (illustrated in Figure 3b) is trained to find a way of representing different environmental conditions such that each species’ response to the environment can be described using a small number of coefficients (e.g. 15 in this analysis; Appendix B). The small number of coefficients and the uniformity of their functions makes mistnet models highly interpretable: the coefficients linking the second hidden layer to a given species’ probability of occurrence essentially describe that species’ responses to a few leading principal components of environmental variation (cf Vincent et al. (2010)). For comparison, the boosted regression tree SDMs used below (Elith, Leathwick & Hastie 2008) have tens of thousands of coefficients per species, with entirely new interpretations for each new species’ coefficients.

How do we train the model to make good predictions? As with most neural networks, mistnet’s coefficients are initialized randomly, and then the model climbs the log-likelihood surface by iteratively adjusting the coefficients toward better values. In mistnet models, the adjustments are calculated with a variant of the backpropagation algorithm (Rumelhart, Hinton & Williams 1986; Murphy 2012) suggested by Tang & Salakhutdinov (2013) for stochastic neural networks. The fitting procedure alternates between inferring the states of the latent variables (via importance sampling) and updating the model’s coefficients (via backpropagation). Both phases of model fitting are described in more detail in Appendix B. Despite importance sampling’s imprecision, this generalized expectation maximization procedure will converge to a local optimum on the likelihood surface with probability one (Neal & Hinton 1998; Tang & Salakhutdinov 2013), ensuring that the expected likelihood is high after averaging over the possible random samples. Following best practices (Orr & Müller 1998; Murphy 2012), mistnet constrains the coefficients using *L*_2_ regularization to prevent overfitting; the strength of this “weight decay” term was chosen by cross-validation, as described in the Appendix.

The mistnet source code can be viewed and downloaded from https://github.com/davharris/mistnet. While the user interface and most of the algorithms are written in R, a small portion of the code is written in C++, using Rcpp (Eddelbuettel & Francois 2011) to manage the interface between languages and RcppArmadillo (Eddelbuettel & Sanderson 2014) to access the Armadillo linear algebra library for faster matrix manipulations (Sanderson 2010).

### Existing models used for comparison

I compared mistnet’s predictive performance with two machine learning techniques and with a linear JSDM called BayesComm (Golding 2013; Golding & Harris 2014). Each of these techniques is described briefly below; implementational details and settings for each method can be found in the Appendix.

The first machine learning method I used for comparison, boosted regression trees (BRT; Elith *et al*. 2008), is among the most powerful techniques available for single-species SDM (Elith *et al*. 2006; Elith *et al*. 2008). I trained one BRT model for each species using R’s `gbm` package (Ridgeway 2013) and stacked them following the recommendations in (Calabrese *et al*. 2014).

I also used a neural network model with no stochastic latent variables as a baseline against which to compare mistnet. Such neural networks do share some information among species (i.e. all species’ log-odds of occurrence are linear combinations of the same hidden layer), but like most other multi-species SDMs (De’ath 2002; Leathwick *et al*. 2005; Ferrier *et al*. 2007) they are not JSDMs and do not explicitly model co-occurrence (Clark *et al*. 2013). The neural net baseline was trained using the `nnet` package (Venables & Ripley 2002).

Finally, I trained a BayesComm model (Golding 2013; Golding & Harris 2014) to evaluate the importance of mistnet’s nonlinearities compared to a linear alternative that also models co-occurrence explicitly.

To ensure a level playing field, each modeling approach was given about 15 hours on the same computer for cross-validation and to make its predictions, as described in the Appendix.

### Evaluating model predictions along test routes

I evaluated mistnet’s predictions both qualitatively and quantitatively. Qualitative assessments involved looking for patterns in the model’s predictions and comparing them with ornithological knowledge (e.g. the habitat classifications provided by the Cornell Lab of Ornithology).

Each model was evaluated quantitatively on the test routes (red points in Figure 2) to assess its predictive accuracy out-of-sample. Models were scored according to their predictive likelihoods, i.e. the probabilities they assigned to various scenarios observed in the test data. Models with high likelihoods expect realistic co-occurrence patterns, and should yield more biologically relevant insights about the processes underlying those patterns. Models that overfit or underfit will have lower out-of-sample likelihoods, and should be trusted less to provide these kinds of insights. I tested each model’s ability to make several kinds of predictions, ranging from estimates of the probability of observing particular species at a given location, to predictions about the species richness and composition of entire assemblages.

To quantify the difficulties each model faced as it made predictions about increasingly large assemblages, I estimated their route-level predictive likelihoods for randomly-chosen groups of species, ranging in size from individual species pairs to the full set of 368 species in the data set. Models that assumed species were uncorrelated should see an exponential decay in their likelihoods as the number of species increases (since the probability of making correct predictions for a set of uncorrelated species equals the product of their individual probabilities), while BayesComm and mistnet should be able to take advantage of correlations to simplify problem of making predictions for the larger assemblages.

Finally, each model predicted a range of possible species richness values for each test route; I calculated quantiles for each model’s predictions using the Poisson-binomial distribution (Hong 2013), as recommended in Calabrese et al. (2014).

## Results and Discussion

### Mistnet’s view of North American bird assemblages

I began by decomposing the variance in the mistnet’s species-level predictions among-routes (which varied in their climate values) and residual variation within routes. On average, the residuals accounted for 29% of the variance in mistnet’s predictions, indicating that non-climate factors play a substantial role in habitat filtering at continental scales.

If the non-climate factors mistnet identified were biologically meaningful, then there should be a strong correspondence between the 15 coefficients assigned to each species by mistnet and the habitat classifications assigned by the Cornell lab of Ornithology. A linear discriminant analysis (LDA; Venables & Ripley 2002) demonstrated such a correspondence (Figure 4). The two-dimensional subspace in Figure 4 explains 19% of the total variance in species’ coefficients (representing an even greater portion of the non-climate variance). Mistnet’s coefficients cleanly distinguished several groups of species by habitat association (e.g. “Grassland” species versus “Forest” species), though the model largely failed to distinguish “Marsh” species from “Lake/Pond” species and “Scrub” species from “Open Woodland” species. These results indicate that the model has identified the broad differences among communities, but that it lacks some fine-scale resolution for distinguishing among types of wetlands and among types of partially-wooded areas. Alternatively, perhaps these finer distinctions are not as salient at the scale of a 40-km transect.

**Figure 4.**
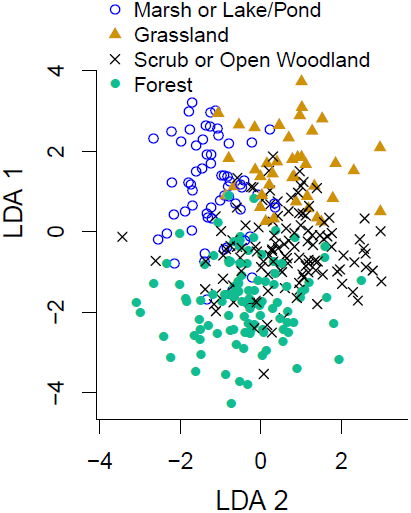
Each species’ mistnet coefficients have been projected into a two-dimensional space by linear discriminant analysis (LDA) in order to maximize the spread between the six habitat types assigned to species by the Cornell Lab of Ornithology’s All About Birds website. The figure shows that mistnet cleanly separates “Grassland” species from “Forest” species, with “Scrub” and “Open Woodland” species representing intermediates along this axis of variation. “Marsh” and “Lake/Pond” species cluster together in the upper-left. The other habitat classes were included in the LDA, but are not shown here.

Figure 5A shows how the forest/grassland gradient identified by mistnet affects the model’s predictions for a pair of species with opposite responses to forest cover. The model cannot tell *which* of these two species will be observed (since it was only provided with climate data), but the model has learned enough about these two species to tell that the probability of observing *both* along the same 40-km transect is much lower than would be expected if the species were uncorrelated.

**Figure 5.**
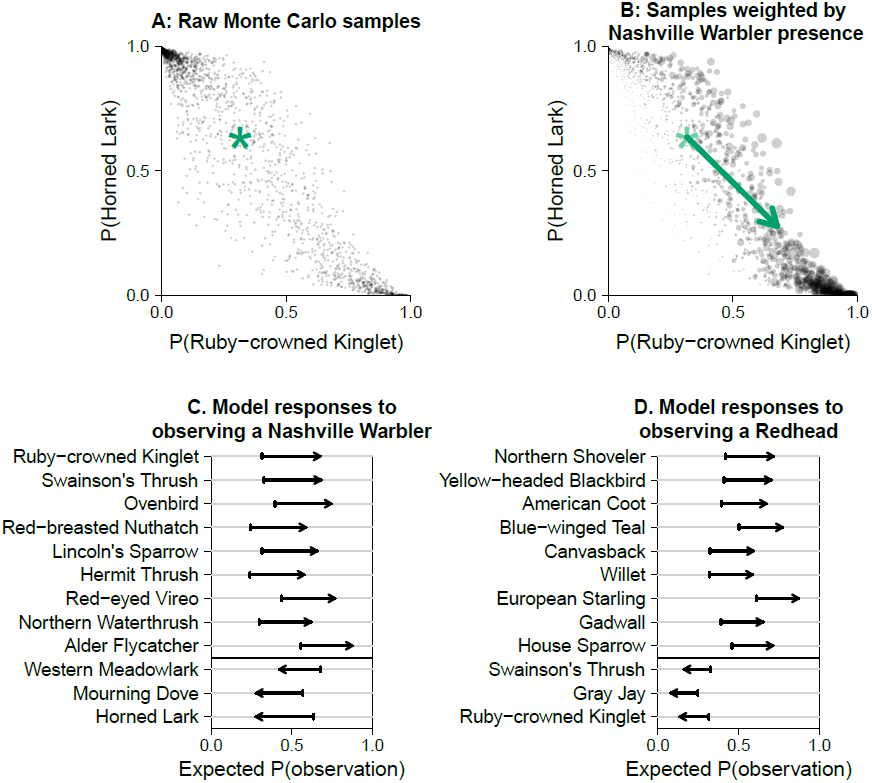
**A**. The mistnet model has learned that Ruby-crowned Kinglets (*Regulus calendula*) and Horned Larks (*Eremophila alpestris*) have opposite responses to some environmental factor whose true value is unknown. Based on these two species’ biology, an ornithologist could infer that this unobserved variable is related to forest cover, with the Kinglet favoring more forested areas and the Lark favoring more open areas. **B**. The presence of a forest-dwelling Nashville Warbler (*Oreothlypis ruficapilla*) provides the model with a strong indication that the area is forested, increasing the weight assigned to Monte Carlo samples that are suitable for the Kinglet and decreasing the weight assigned to samples that are suitable for the lark. **C**. The Nashville Warbler’s presence similarly suggests increased occurrence probabilities for a variety of other forest species, as well as decreased probabilities for species associated with wetlands and grasslands. **D**. If a Redhead (*Aythya americana*) has been observed along a route, the model correctly expects to see more ducks, rails and sandpipers in the same area.

Figure 5A reflects a great deal of uncertainty, which is appropriate considering that the model has no information about a crucial environmental variable (forest cover). Often, however, additional information is available that could help resolve this uncertainty, and the mistnet package includes a built-in way to do so, as indicated in Figures 5B and 5C. These panels show how the model is able to use an observation of a forest-associated Nashville Warbler (*Oreothlypis ruficapilla*) to indicate that a whole suite of other forest-dwelling species are likely to occur nearby, and that a variety of species that prefer open fields and wetlands should be absent. Similarly, Figure 5D shows how the presence of a Redhead duck (*Aythya americana*) can inform the model that a route is suitable habitat for a variety of other ducks, as well as for other wetland-associated species such as marsh-breeding blackbirds, sandpipers, and rails (along with a few other species that do not fit this theme as nicely). None of these inferences would be possible from a stack of disconnected single-species SDMs.

### Model comparison: species richness

Environmental heterogeneity plays an especially important role in determining species richness, which is often overdispersed relative to models’ expectations (O’Hara 2005). Figure 6 shows that mistnet’s predictions respect the heterogeneity one might find in nature: areas with a given climate could be largely unsuitable for waterfowl (Anatid richness < 2 species) or marshy and open (Anatid richness > 10 species). Under the independence assumption used for stacking SDMs, however, both of these very plausible scenarios would be ruled out (Figure 6A).

**Figure 6.**
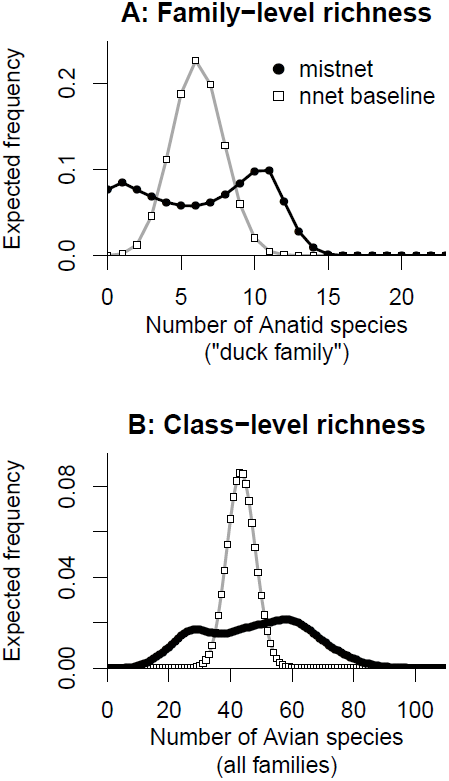
The predicted distribution of species richness one would expect to find based on predictions from mistnet and the baseline neural network. A. Anatid species (waterfowl). B. All bird species. BRT’s predictions (not shown) are similar to the baseline network, since neither one accounts for the effects of unmeasured environmental heterogeneity.

Unfortunately, stacking leads to even larger errors when predicting richness for larger groups, such as the complete set of birds studied here. Models that stacked independent predictions underestimated the range of biologically possible outcomes (Figure 6B), frequently putting million-to-one or even billion-to-one odds against species richness values that were actually observed. In more concrete terms, half of the observed species richness values fell outside these models’ 95% confidence intervals. The overconfidence associated with stacked models could have serious consequences in both management and research contexts if we fail to prepare for species richness values outside such an unreasonably narrow range.

Mistnet, on the other hand, was able to explore the range of possible non-climate environments to avoid these missteps: 90% of the test routes fell within mistnet’s 95% confidence intervals, and the log-likelihood ratio decisively favored it over stacked alternatives.

### Model comparison: single species

The two neural network models had the best performance at the level of individual species (Table 1). The neural networks’ advantage over BRT was largest for low-prevalence species (linear regression of log-likelihood ratio versus log-prevalence; p = 0.004). This is consistent with previous observations that multi-species models can outperform single-species approaches for rare species (Leathwick, Elith & Hastie 2006), which will often be of the greatest conservation concern. BayesComm’s predictions were substantially worse than any of the machine learning methods, which I attribute to its inability to learn nonlinear responses to the environment.

**Table 1.** Expected species-level log-likelihood for each method, summed over all test routes and averaged across all species. The likelihood ratio compares each model to BRT, representing single-species SDMs. Sharing information among species with either of the neural net models improves the predictive likelihood more than twenty-fold for a typical species compared to BRT. Note also that BayesComm averages less than 1% of the machine learning methods’ likelihoods because of its linearity assumption.

| method | expected.log.likelihood | likelihood.ratio |
| --- | --- | --- |
| nnet | -48.7 | 21.3 |
| mistnet | -48.7 | 21 |
| BRT | -51.7 | 1 |
| BayesComm | -56.6 | 0.00771 |

### Model comparison: community composition

While making predictions about individual species observations is fairly straightforward with this data set (since most species have relatively narrow breeding ranges), community ecology is more concerned with co-occurrence and related patterns involving community composition (Chase 2003). As expected, models that combined their single-species predictions independently (including the neural network baseline) showed exponential decay in their likelihoods as the number of species per prediction increased. The JSDMs (mistnet and BayesComm) showed sub-exponential declines, since correlations reduce the number of independent bits of information needed to make an accurate prediction. As a result, mistnet became increasingly advantageous over independent combinations of single-species predictions as the assemblage size increased (Figure 7). Mistnet’s log-likelihood averaged 10.8 units higher than BRT’s for full assemblages of 368 species, corresponding to a 47000-fold improvement in likelihood for a typical transect in the test set. Mistnet’s ability to focus its predictions on plausible combinations of species indicates that it has captured a great deal more of the underlying ecological processes than existing SDM approaches. While some of this improvement can be attributed to mistnet’s overall tendency to make better predictions about individual species (Table 1), the difference is mainly due to mistnet’s ability to keep ahead of the combinatorial explosion of possible assemblages by exploiting correlations among species.

**Figure 7.**
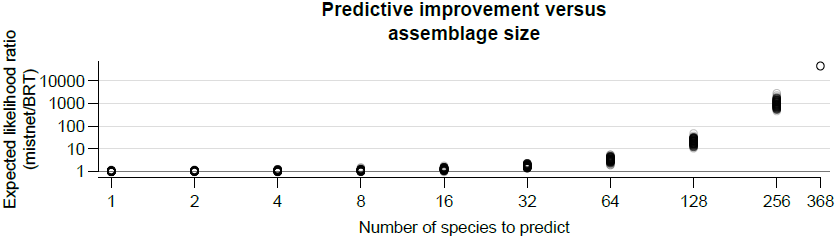
The likelihood ratio favoring mistnet over BRT grows super-exponentially with assemblage size. Each circle corresponds to a randomly-generated set of *N* species, where the value of *N* is indicated along the horizontal axis. Note the log scale on both axes.

### Comparison with BayesComm

BayesComm’s ability to make out-of-sample predictions was severely limited by its assumption that species respond linearly to climate variables, highlighting the the need for nonlinear methods that can learn the functional forms of species’ responses to the environment. Adding quadratic and interaction terms would have led to severe overfitting for many rare species, and may still not have provided enough flexibility to compete with nonlinear techniques.

Even without the added complexity of nonlinear terms, the BayesComm model required 70,000 parameters, most of which served to to identify a distinct correlation coefficient between a single pair of species. Tracing this many parameters through hundreds of Markov chain iterations routinely caused BayesComm to exceed my machine’s 8 gigabytes of memory and crash, even after the code was modified to reduce its memory footprint. Storing long Markov chains over a dense, full-rank covariance matrix (as has apparently been done in all other JSDMs to date) thus appears not to be a feasible strategy with large assemblages.

## Conclusion

These results show conclusively that both linearity and independence are unwarranted assumptions; either assumption can substantially impair our ability to model and understand large assemblages. Linear JSDMs are not flexible enough, and models without latent random variables cannot match the properties of real assemblages.

SDMs’ failure to sufficiently consider correlations among has kept these models from explaining and anticipating the full range of complex assemblages found in nature (Austin & Van Niel 2011). Mistnet’s predictions are much more compatible with these sorts of complexities. In particular, the model’s predictions need not be unimodal, allowing the model to express conditional predictions, such as that “the probability of observing a Redhead duck will be very high if other wetland species are present, but very low otherwise.” Such conditional predictions are important because the available data will not always contain enough information to narrow the possibilities down to a single assemblage type or a single group of species. In such situations, stacked models will provide a false sense of security out-of-sample, leading to bad decisionmaking and biased estimates of nature’s variability. Mistnet provides better confidence intervals that are much more likely to actually contain the observed values when we look out-of-sample.

Mistnet can also identify some of the same similarities among species that a skilled biologist would expect to find, which will be important for studying taxa that are more diverse and harder to observe (such as microbes). For taxa on the frontier of our knowledge, a model like mistnet could help guide the biologists to ask the best questions and organize their understanding by suggesting which species have similar habitat requirements—even when the factor controlling their occurrence are still unknown.

Unlike with stacked methods, one can read this straight out of mistnet’s coefficient tables with no more difficulty than interpreting a Principal Components Analysis.

Mistnet’s ability to use asymmetrical or low-quality data sources to improve its predictions should inrease the value of low-effort data collection procedures such as short transects— especially since these improvements can be incorporated without need for fitting a new model. Future research should look for ways to use other forms of ecological knowledge about species to impose some structure on models coefficients and nudge the models toward more biologically reasonable predictions (Kearney & Porter 2009; Kissling *et al*. 2012). Such a research program could also be useful in other areas of predictive ecology [@pearse predicting 2013].

Finally, it should be noted that, while one *could* describe direct interactions among species using latent variables (Ovaskainen *et al*. 2010; Golding 2013), existing JSDMs are not particularly well-suited for learning about species interactions. Other models, such as Markov random fields (Azaele *et al*. 2010), or ensembles of classifier chains (Yu *et al*. 2011) would be much more appropriate for inferring coefficients related to species interactions, as they include direct dependencies among species. Latent variable-based JSDMs, including mistnet, are more appropriate for studies like this one at large spatial scales where direct species interactions will tend to be weaker and most of the variation is driven by environmental filtering and species’ range limits.

In conclusion, mistnet’s accuracy, as well as its flexibility to work with opportunistic samples should make it useful for a variety of basic and applied contexts. Assemblage-level models, such as mistnet, also have the potential to yield new biological insights. With charismatic and well-studied species like North American birds, most models will mainly be telling information that we already know. Still, mistnet’s ability to capture useful information about axes of variation among birds and to match preconceptions about which species co-occur due to habitat variables may indicate that the model can teach us new things about taxa that are harder to study.

## Acknowledgements

This work benefitted greatly from discussions with A. Sih and his lab meeting group, M. L. Baskett, R. J. Hijmans, R. McElreath, J. H. Thorne, M. W. Schwartz, B. M. Bolker, R. E. Snyder, A. C. Perry, and C. S. Tysor. It was funded by a Graduate Research Fellowship from the National Science Foundation, the UC Davis Center for Population Biology, and the California Department of Water Resources. I gratefully acknowledge the field biologists that collected the BBS data, as well as the US Geological Survey, Worldclim, and Cornell Lab of Ornithology for their efforts and for making their data sets publicly available.

## Data Accessibility

- All data sets used here are freely downloadable from their original sources.
- The mistnet source code can be downloaded from https://github.com/davharris/mistnet/. The easiest way to install the package is with the `devtools` package’s `install_github` command (e.g. `devtools::install_github(“mistnet”, “davharris”`).
- Some code has been improved since the analyses were run; however, the web site includes a complete version history. The analyses in this paper had essentially all been run by the commit at https://github.com/davharris/mistnet/tree/1e2eaaeabf9b4b4360f19b00c0d06508578d7f15.

